# NetTIME: a Multitask and Base-pair Resolution Framework for Improved Transcription Factor Binding Site Prediction

**DOI:** 10.1101/2021.05.29.446316

**Authors:** Ren Yi, Kyunghyun Cho, Richard Bonneau

## Abstract

**Motivation:** Machine learning models for predicting cell-type-specific transcription factor (TF) binding sites have become increasingly more accurate thanks to the increased availability of next-generation sequencing data and more standardized model evaluation criteria. However, knowledge transfer from data-rich to data-limited TFs and cell types remains crucial for improving TF binding prediction models because available binding labels are highly skewed towards a small collection of TFs and cell types. Transfer prediction of TF binding sites can potentially benefit from a multitask learning approach; however, existing methods typically use shallow single-task models to generate low-resolution predictions. Here we propose NetTIME, a multitask learning framework for predicting cell-type-specific transcription factor binding sites with base-pair resolution.

**Results:** We show that the multitask learning strategy for TF binding prediction is more efficient than the single-task approach due to the increased data availability. NetTIME trains high-dimensional embedding vectors to distinguish TF and cell-type identities. We show that this approach is critical for the success of the multitask learning strategy and allows our model to make accurate transfer predictions within and beyond the training panels of TFs and cell types. We additionally train a linear-chain conditional random field (CRF) to classify binding predictions and show that this CRF eliminates the need for setting a probability threshold and reduces classification noise. We compare our method’s predictive performance with two state-of-the-art methods, Catchitt and Leopard, and show that our method outperforms previous methods under both supervised and transfer learning settings.

**Availability:** NetTIME is freely available at https://github.com/ryi06/NetTIME and the code is also archived at https://doi.org/10.5281/zenodo.6994897

## 1. Introduction

Genome-wide modeling of non-coding DNA sequence function is among the most fundamental and yet challenging tasks in biology. Transcriptional regulation is orchestrated by transcription factors (TFs), whose binding to DNA initiates a series of signaling cascades that ultimately determine both the rate of transcription of their target genes and the underlying DNA functions. Both the cell-type-specific chromatin state and the DNA sequence affect the interactions between TFs and DNA *in vivo* [1]. Experimentally determining cell-type-specific TF binding sites is made possible through high-throughput techniques such as chromatin immunoprecipitation followed by sequencing (ChIP-seq) [2]. Due to experimental limitations, however, it is infeasible to perform ChIP-seq (or related single-TF-focused experiments) on all TFs across all cell types and organisms [1]. Therefore, computational methods for accurately predicting *in vivo* TF binding sites are essential for studying TF functions, especially for less well-known TFs and cell types.

Multiple community crowdsourcing challenges have been organized by the DREAM Consortium^1^ to find the best computational methods for predicting TF binding sites in both *in vitro* and *in vivo* settings [3, 4]. These challenges also set the community standard for both processing data and evaluating methods. However, top-performing methods from these challenges have revealed key limitations in the current TF binding prediction community. Generalizing predictions beyond the training panels of cell types and TFs can potentially benefit from multitask learning and increased prediction resolution. However, many existing methods still use shallow single-task models. Predictions generated from these methods typically have low resolution, and they cannot achieve competitive performance for prediction of binding regions shorter than 50 base pairs (bp), although the actual TF binding sites are considerably shorter [5].

### 1.1. Related work

Early TF binding prediction methods such as MEME [6, 7] focused on deriving interpretable TF motif position weight matrices (PWMs) that characterize TF sequence specificity. Amid rapid advancement in machine learning, however, growing evidence has suggested that sequence specificity can be more accurately captured through more abstract feature extraction techniques. For example, a method called DeepBind [8] used a convolutional neural network to extract TF binding patterns from DNA sequences. Several modifications to Deep-Bind subsequently improved model architecture [9] as well as prediction resolution [10]. Yuan et al. developed BindSpace, which embeds TF-bound sequences into a common high-dimensional space. Embedding methods like BindSpace belong to a class of representation learning techniques commonly used in natural language processing [12] and genomics [13, 14] for mapping entities to vectors of real numbers. New methods also explicitly model protein binding sites with multiple binding mode predictors [15], and the effect of sequence variants on non-coding DNA functions at scale [16, 17, 18]. In general, the DNA sequence at a potential TF binding site is just the beginning of the full DNA-function picture, and the state of the surrounding chromosome, the TF and TF-cofactor expression, and other contextual factors play an equally large role. This multitude of factors changes substantially from cell type to cell type. *In vivo* TF binding site prediction, therefore, requires cell-type-specific data such as chromatin accessibility and histone modifications. Convolutional neural networks as well as TF- and cell-type-specific embedding vectors have both been used to learn cell-type-specific TF binding profiles from DNA sequences and DNase-seq data [19]. The DREAM Consortium also initiated the ENCODE-DREAM challenge to systematically evaluate methods for predicting *in vivo* TF binding sites [4]. Apart from carefully designed model architectures, top-ranking methods in this challenge also rely on extensively curated feature sets. One such method, called Catchitt [20], achieves top performance by leveraging a wide range of features including DNA sequences, genome annotations, TF motifs, DNase-seq, and RNA-seq.

### 1.2. Current limitations

Compendium databases such as ENCODE [21] and Remap [22] have collected ChIP-seq data for a large collection of TFs in a handful of well-studied cell types and organisms [1]. Within a single organism, however, the ENCODE TF ChIP-seq collection is highly skewed towards only a few TFs in a small collection of well-characterized cell lines and primary cell types (Figure S1). Transfer learning from well-known cell types and TFs is crucial for understanding less-studied cell types and TFs. One way to achieve transfer learning is by reusing information from a previously learned task to improve learning efficiency of a related task [23]. For example, pretraining machine learning models with data from multiple TFs allows the models to learn common binding characteristics among TFs and thus, improves fine-tuning performance on a single TF of interest [24, 25]. Existing methods that adopt the above transfer learning approach [24, 25] do not yet include model components that account for the TF and cell type identities in an integrated fashion, which makes the fine-tuning step necessary for predicting binding preferences for a particular TF of interest. In contrast, multitask learning models that can account for TF and cell type identities eliminate the necessity of fine-tuning when learning to predict binding preferences for new TFs in new cell types (this in turn enables a more meaningful integration of much larger training sets). Moreover, as different TFs have different binding mechanisms under various cellular conditions [26], models that can account for TF and cell type identities are potentially more effective at transfer learning compared to models, such as [24] and [25], that do not have proper strategy for recognizing binding data of different TF and cell type origins.

Top-performing methods from the ENCODE-DREAM Challenge typically adopt the single-task learning approach. For example, Catchitt [20] trains one model per TF and cell type. Cross cell-type transfer predictions are achieved by providing a trained model with input features from a new cell type. This approach can be highly unreliable as the chromatin landscapes between the trained and predicted cell types can be drastically different [27] and these differences can be functionally important [28]. Alternatively, each model can be trained on one TF using cell-type-specific data across multiple cell types of interest [29]. Without proper mechanisms to distinguish cell types, however, such models tend to assign high binding probabilities to common binding sites among training cell types. A few methods have adopted the multitask learning approach in which data from multiple cell types and TFs are trained jointly in order to improve the overall model performance [16, 30, 17, 31, 18, 32, 33]. The multitask solution adopted by DeepSea [16] and several other methods [30, 17, 31, 18, 32] involves training a multiclass classifier on DNA sequences, where each class represents the occurrence of binding sites for one TF in one cell type. This solution is suboptimal as it cannot generalize predictions beyond the training TF and cell-type pairs. Sequence context affects TF binding affinity [34], and increasing context size can improve TF binding site prediction [16]. TF binding sites are typically only 4-20 bp long [5]; TF binding models that can achieve base-pair prediction resolution are therefore beneficial for experimental validation as well as *de novo* motif discovery. However, instead of identifying precise TF binding locations, existing methods mainly focus on determining the presence of binding sites. Predictions from these models suffer from either low resolution or low context size, depending on the input sequence length. Leopard [35] and BPNet [33] are two recently proposed base-pair resolution binding prediction methods for predicting cell-type-specific TF binding sites. Leopard uses both DNA sequences and DNase-seq chromatin accessibility data as input, whereas BPNet predicts binding sites solely from DNA sequences. However, Leopard is a single-task learning model that requires training one model per TF and per cell type. Although BPNet uses multitask learning, the model does not include any task-specific components for distinguishing different TF and cell type identities, and it’s performance when training on more than four conditions (described in [33]) has not been evaluated.

In this work, we address the above challenges by introducing NetTIME (Network for TF binding Inference with Multitask-based condition Embeddings), a multitask learning framework for base-pair resolution prediction of cell-type-specific TF binding sites. NetTIME jointly trains multiple cell types and TFs, and effectively distinguishes different conditions using cell-type-specific and TF-specific embedding vectors. It achieves base-pair resolution and accepts input sequences up to 1kb.

## 2. Approach

### 2.1. Feature and label generation

The ENCODE Consortium has published a large collection of TF ChIP-seq data, all of which are generated and processed using the same standardized pipelines [21]. We therefore collect our TF binding target labels from ENCODE to minimize data heterogeneity. Each replicated ENCODE ChIP-seq experiment has two biological replicates, from which two sets of peaks – conserved and relaxed – are derived; peaks in both sets are highly reproducible between replicates^2^. Compared to the relaxed peak set, the conserved peak set is generated with a more stringent threshold, and is generally used to provide target labels. However, the conserved peak set usually contains too few peaks to train the model efficiently. Therefore, we use both conserved and relaxed peak sets to provide target labels for training, and the conserved peak set alone for evaluating model performance.

To collect target labels for a representative set of conditions that cover a wide range of cellular conditions and binding patterns, we first select 7 cell types and 22 TFs for which ENCODE has available binding data. The 7 cell types include 3 cancer cell types, 3 normal cell types and 1 stem cell type. The 22 TFs include 17 TFs from 7 TF protein families as well as 5 functionally related TFs. Conserved and relaxed peak sets are collected from 71 ENCODE replicated ChIP-seq experiments conducted on our cell types and TFs of interest. Each of these TF ChIP-seq experiment is henceforth referred to as a condition. All peaks from these conditions form a set of information-rich regions where at least one TF of interest is bound. We generate samples by selecting non-overlapping *L*-bp genomic windows from this information-rich set, where *L* is the context size. We set the context size *L* = 1000 as it was previously shown to improve TF binding prediction performance [16].

*In vivo* TF binding sites are affected by DNA sequences and the cell-type-specific chromatin landscapes. In addition to using DNase-seq, which maps chromatin accessibility, we collect ChIP-seq data for 3 types of histone modifications to form our cell-type-specific feature set. The histone modifications we include are H3K4me1, H3K4me3 and H3K27ac, which are often associated with enhancers [36], promoters [37] and active enhancers [38], respectively.

### 2.2. Methods

NetTIME performs TF binding predictions in three steps: 1) generating the feature vector **w** = (*w*_1_, …, *w*_*L*_) given a TF label *p*, a cell type label *q*, and a sample DNA sequence **x** = (*x*_1_, …, *x*_*L*_) where each *x*_*l*_ ∈ {*A, C, G, T*}, 2) training a neural network to predict base-pair resolution binding probabilities **z** = (*z*_1_, …, *z*_*L*_), and 3) converting binding probabilities to binary binding decisions **y** = (*y*_1_, …, *y*_*L*_) of *p* in *q* by either setting a probability threshold or additionally training a conditional random field (CRF) classifier (Figure 1).

**Figure 1:**
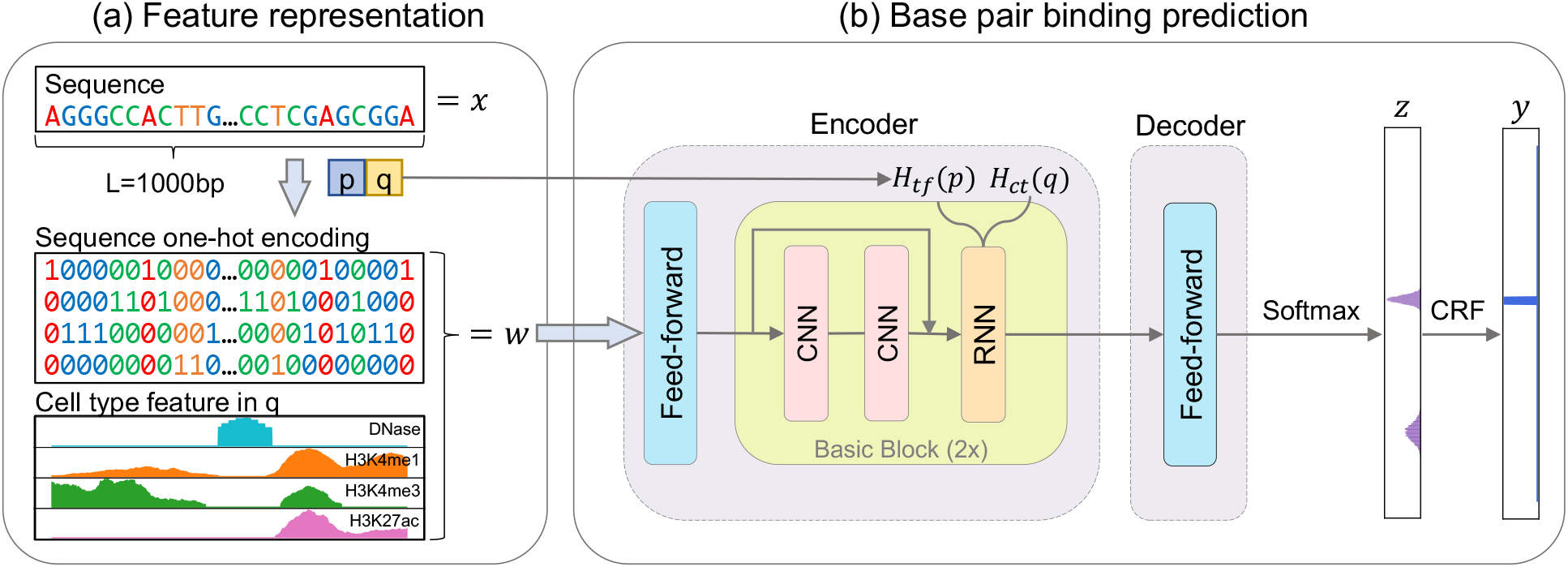
Schematic method overview. **(a)** Constructing feature vector *w* from input sequence *x*, TF label *p* and cell type label *q. w* consists of the sequence one-hot encoding, and a set of cell-type-specific features – DNase-seq signals, and H3K4me1, H3K4me3 and H3K4ac histone ChIP-seq signals – in cell type *q*. **(b)** Feature vector *w*, TF label *p* and cell type label *q* are provided to the NetTIME neural network to predict base-pair resolution binding probability *z*. An additional CRF classifier is trained to predict binary binding event *y* from *z*.

#### 2.2.1. Feature representation

We construct the feature vector **w** ∈ ℝ^*K*×*L*^ from **x** ∈ ℝ^*L*^, where *K* represents the number of features. Different types of features are independently stacked along the first dimension. For each element in **w**, *w*_*l*_ is the concatenation of the one-hot encoding of the DNA sequence *O*(*x*_*l*_), and the cell-typespecific feature *C*(*x*_*l*_) (Figure 1a).

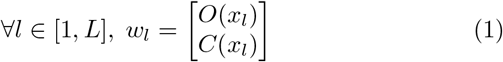

High-dimensional embedding vectors can be trained to distinguish different conditions as well as implicitly learning condition-specific features, and are therefore preferred by many machine learning models over one-dimensional condition labels [14, 19, 11]. Given TF label *p* and cell type label *q*, NetTIME learns the TF- and cell-type-specific embeddings *H*_*tf*_ (*p*) ∈ ℝ^*d*^ and *H*_*ct*_(*q*) ∈ ℝ^*d*^*′*, where *d* = *d*′ = 50.

#### 2.2.2. Binding probability prediction

NetTIME adopts an encoder-decoder structure similar to that of neural machine translation models [39, 40, 41] (Figures 1b, S2):

*Encoder:*. the model encoder maps the input feature **w** to a hidden vector **h** ∈ ℝ^2*d*×*L*^. The main structure of the encoder, called the Basic Block, consists of a CNN followed by a recurrent neural network (RNN). CNN uses multiple short convolution kernels to extract local binding motifs, whereas bi-directional RNN is effective at capturing longrange TF-DNA interactions [42, 43]. We choose the ResBlock structure introduced by ResNet [44] as our CNN, as it has become a standard approach for training deep neural networks [41, 45]. Traditional RNNs are challenging to train due to the vanishing gradient problem [42]. We therefore use the bi-directional gated recurrent unit (bi-GRU) [43], a variant of RNN proposed to address the above challenge. The hidden state of bi-GRU is initialized by concatenating the embedding vectors *H*_*tf*_ (*t*) and *H*_*ct*_(*c*).

*Decoder:*. the model decoder converts the hidden vector **h** to binding probabilities **z**. The conversion is achieved through a fully connected feed-forward network, as the relationship between **h** and **z** may not be trivial. A softmax function subsequently transforms the decoder output to the binding probabilities.

#### 2.2.3. Training

We train the model by minimizing the negative conditional log-likelihood of **z**:

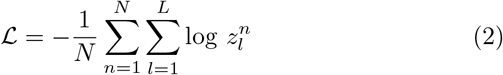

where *N* denotes the number of training samples. The loss function is optimized by the Adam optimizer [46] (also see Supplementary Section 1.2).

#### 2.2.4. Binding event classification

Binary binding events **y** can be directly derived from **z** by setting a probability threshold *b* ∈ (0, 1) such that

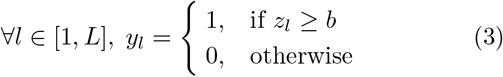

This approach has been used by many exising TF binding predictions models to admit exact inference [35, 47]. Alternatively, a linear-chain CRF classifier can be trained to achieve the same goal. It computes the conditional probability of **y** given **z**, defined as the following:

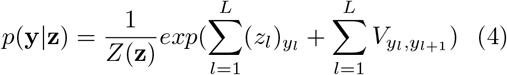

1. *Z*(**z**) is a normalization factor,
2. *V* ∈ *ℝ*^*p*×*p*^ is a transition matrix, where *p* denotes the number of classes of the classification problem and each *V*_*i,j*_ represents the transition probability from class label i to j,
3. 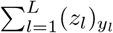 calculates the likelihood of *y*_*l*_ given *z*_*l*_, and
4. 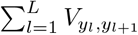 measures the likelihood of given *y*_*l*_.

In CRF, the class label at position *l* affects the classification at position *l* + 1 [48]. This is potentially beneficial for TF binding site classification as positions adjacent to a putative binding site are also highly likely to be occupied by TFs. We train the CRF by minimizing −log *p*(**y**|**z**) over all training samples. The Adam optimizer [46] is used to update the parameter *V*.

### 2.3. Model selection

We follow the guideline provided by the ENCODE-DREAM Challenge [4] to perform data split as well as model selection whenever possible. Training, validation and test data are split according to chromosomes (Supplementary Table S1). We use the area under the Precision-Recall Curve (auPRC) score to select the best neural network model checkpoint.

To access how well our model predictions recover the positive binding sites in the truth target labels, we evaluate classifiers’ performance according to Intersection Over Union (IOU) score. Suppose *P* and *T* are sets of predicted and target binding sites, respectively. Then

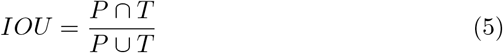

We test 300 random probability thresholds and select the best threshold, i.e., the threshold that achieves the highest IOU score in the validation set. We also train a CRF using predictions generated from the best neural network checkpoint. The best CRF checkpoint is selected according to the average loss on the validation set. Model performance reported here is evaluated using the test set.

## 3. Results

### 3.1. Multitask learning improves performance by increasing data availability

NetTIME can be trained using data from a single condition (single-task learning) or multiple conditions (multitask learning). Jointly training multiple conditions allows the model to use data more efficiently and improves model generalization [49]. Multitask learning is particularly suitable for learning cell-type-specific TF binding preferences because a TF has common binding sites across different cell types, and functionally related TFs share similar binding sites [50]. We therefore evaluate the effectiveness of multitask learning when jointly training multiple related conditions. For this analysis, we choose three TFs from the JUN family that exhibit overlapping functions: JUN, JUNB, and JUND [51]. Combining multiple cell types of JUND allows the multitask learning model to significantly outperform the single-task learning models, each of which is trained with one JUND condition (Figure 2a). Jointly training multiple JUN family TFs further improves performance compared to training each JUN family TF separately (Figure 2b). However, we observe decreased performance when subsampling the multitask models’ training data to match the number of samples in the corresponding single-task models (Figure 2).

**Figure 2:**
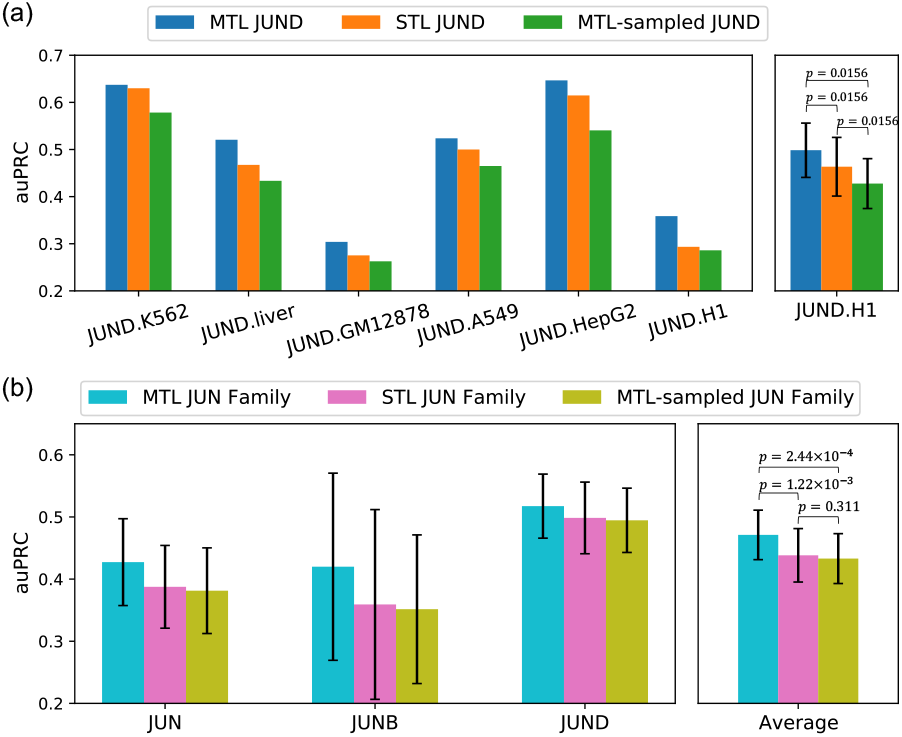
Performance comparison between multitask learning and single-task learning approaches using JUN family TFs. Models are trained with datasets from JUND across multiple cell types, and **(b)** multiple TFs in the JUN family across multiple cell types. MTL: multitask learning; MTL-sampled: multitask learning training data that has been subsampled to match the number of samples in the corresponding single-task models; STL: single-task learning. The right panels in (a) and (b) are the averaged auPRC of the models shown in the corresponding left panels. Error bars represent standard error of the mean across all training conditions. P-values are calculated using the Wilcoxon signed-rank test using auPRC scores across all conditions.

This indicates that the multitask learning strategy is more efficient due to the increased data available to the multitask models rather than to the increased data diversity. Similar results are also observed when the same analysis is performed on three unrelated TFs (Figure S3).

### 3.2. Supervised predictions made by NetTIME outperforms existing baseline methods

Our complete feature set includes DNA sequence, and cell-type-specific features including DNase-seq and three types of histone ChIP-seq. In practice, however, data for these features are not always available for the conditions of interest. Additionally, TF motif enrichment has often been used by existing methods to provide TF binding sequence specificity information [20, 29]. We therefore evaluate the quality of our model predictions when we vary the types of input features available during training.

We first train separate models after removing cell-type-specific features using training data from all conditions mentioned in Section 2.1. Model prediction accuracy is evaluated in the supervised fashion using the test data from the same set of conditions. The addition of cell-type-specific features significantly improves NetTIME performance. However, adding TF motif enrichment features (Supplementary Section 1.1), either in addition to DNA sequence features or in addition to both sequence and cell type features, does not introduce significant performance improvement (Figure 3a). Despite exhibiting high sequence specificity *in vitro*, TF binding sites *in vivo* correlate poorly with TF motif enrichment [52]. Motif qualities in TF motif databases vary significantly depending on the available binding data and motif search algorithms. Nevertheless, TF motifs have been the gold standard for TF binding site analyses due to their interpretability and scale. However, TF motif enrichment features are likely redundant when our model can effectively capture TF binding sequence specificity, though it’s possible our protocol for generating TF motif enrichment features is suboptimal.

**Figure 3:**
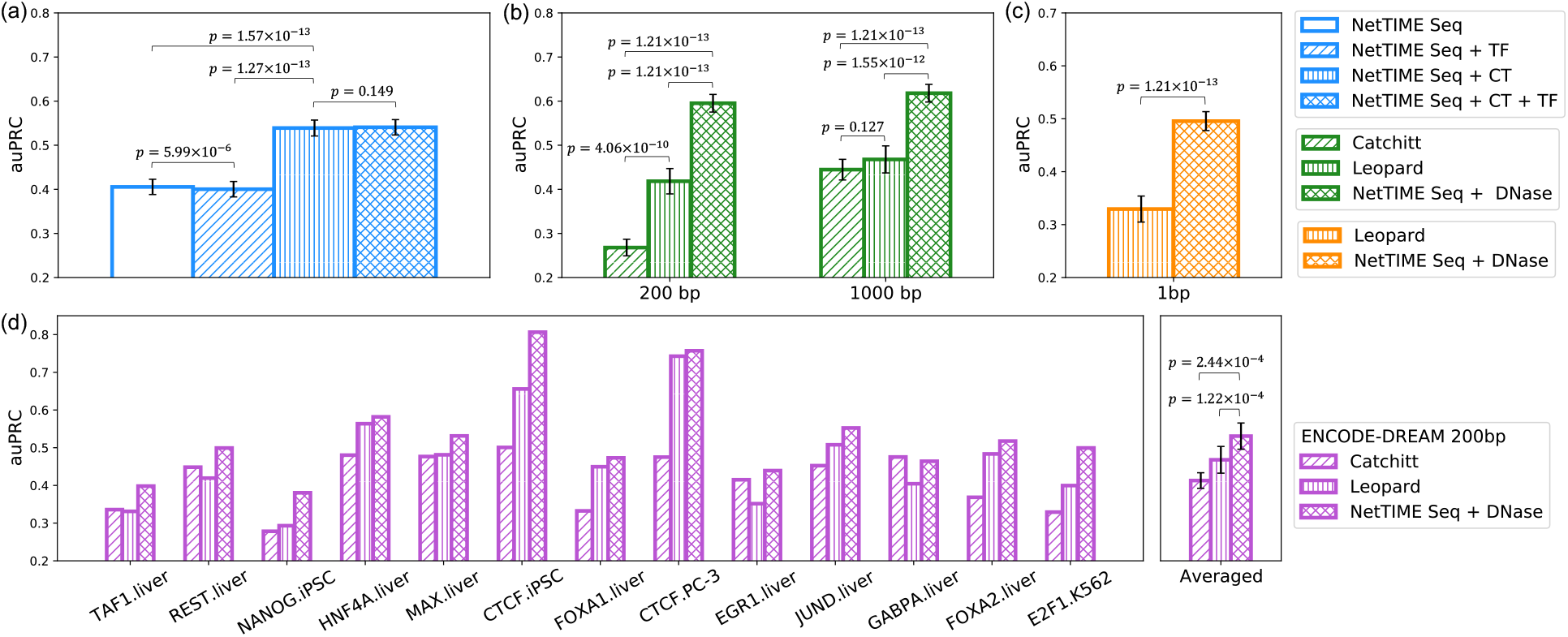
NetTIME significantly improves predictive performance against state-of-the-art baseline methods. **(a)** Comparing NetTIME performance under different input feature settings. Seq: DNA sequence feature; CT: cell-type-specific features including DNase-seq, and H3K4me1, H3K4me3 and H3K27ac histone ChIP-seq data; TF: TF motif enrichment feature. **(b)** Comparing NetTIME performance **(b)** against Catchitt and Leopard under 200-bp and 1000-bp resolutions, and **(c)** against Leopard under 1-bp resolution. DNase: cell-type-specific DNase-seq feature. **(d)** Comparing NetTIME Seq + DNase model performance against Catchitt and Leopard under 200-bp resolution using the ENCODE-DREAM Challenge data.

We further compare NetTIME predictive performance with that of Catchitt [20] and Leopard [35]. As Catchitt and Leopard use only DNase-seq data as their cell-type-specific input feature, we train a separate NetTIME model using DNA sequences and DNase-seq data as input. Additionally, because Catchitt is evaluated under the 200-bp resolution for the ENCODE-DREAM Challenge, we reduce the NetTIME and Leopard prediction resolution by taking the maximum prediction probability across the center 200-bp regions for all the example sequences in our test set. Performance of these three methods are further compared under 1000-bp resolution to evaluate per-sample prediction accuracy. Prediction auPRC scores consistently increase for all three methods as we decrease the prediction resolution from 200-bp to 1000-bp. Nevertheless, NetTIME outperforms both baseline methods under both prediction resolutions (Figure 3b). Furthermore, NetTIME significantly outperforms Leopard when predictions are evaluated on the per-basepair level (Figure 3c). Although the TF motifs are not used by Leopard as a type of input feature, Leopard derives target binding labels by subsetting TF ChIP-seq peaks with regions that show TF motif enrichment [35]. This data generation procedure potentially introduces unwanted biases and contributes to the reduced performance when the model is evaluated on the complete set of TF ChIP-seq peaks.

Both Catchitt and Leopard can only be trained using examples derived genome-wide. To ensure a fair comparison, we train additional Seq + DNase NetTIME models using DNase seq data and ChIP-seq labels provided by the ENCODE-DREAM Challenge. All three methods are benchmarked against the 13 test conditions in the ENCODE-DREAM Challenge, and their model performance is evaluated at 200-bp resolution using examples generated by sliding a 200-bp window across all test chromosomes with a 50-bp overlap between adjacent examples. Predictions at 200-bp resolution from NetTIME and Leopard are generated by taking the maximum probability across each 200-bp region from the 1-bp resolution predictions generated by these two methods. NetTIME improves the mean prediction auPRC score by 11.8% and 6.3% over Catchitt and Leopard, respectively (Figure 3d).

### 3.3. TF- and cell-type-specific embeddings are crucial for an effective multitask learning strategy

Here we evaluate the relative contributions of different model components to our predictive accuracy. We use the TF and cell-type embedding vectors to learn condition-specific features and biases, and a combination of CNNs and RNNs to learn the non-condition-specific TF-DNA interaction patterns. TF and cell type embedding vectors can be replaced with random vectors at prediction time and at training time to evaluate the contribution of each component individually.

To evaluate the model’s sensitivity to different TF and cell type labels, TF and cell type embedding vectors are replaced with random vectors at prediction time (Figure 4). When NetTIME is trained with both TF and cell type embeddings, the model learns to use both pieces of condition-specific information in order to make accurate predictions. As a result, substituting both types of embeddings with random vectors reduces our model performance by 69.1% on average. Replacing either TF or cell type embeddings with random vectors also drastically reduce auPRC scores. This indicates that cell-type-specific chromatin landscape, in addition to TF identity, is important for defining *in vivo* TF binding sites, which explains the redundancy of TF motif features and the lack of correlation between TF ChIP-seq signals and TF motif enrichment mentioned in Section 3.2 and [52].

**Figure 4:**
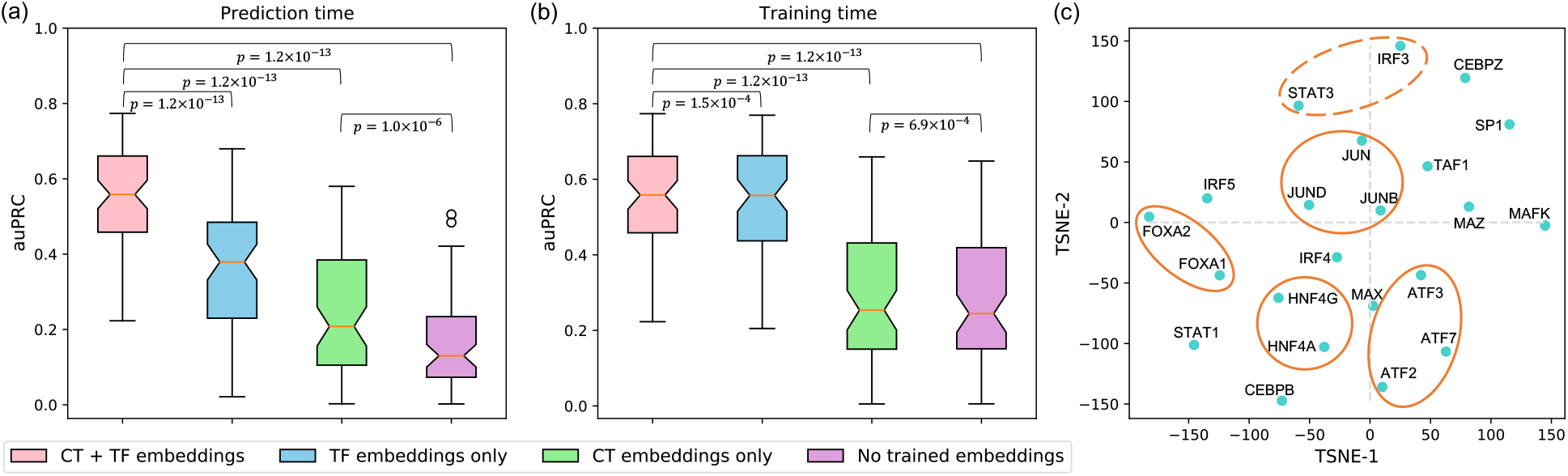
Properties of trained embedding vectors. Evaluating the contribution of condition-specific network components **(a)** at prediction time and **(b)** at training time by replacing trained TF and cell type embeddings with random vectors. CT + TF embeddings: trained TF and cell type embeddings; TF embeddings only: trained TF embeddings and random cell type vectors; CT embeddings only: trained cell type embeddings and random TF vectors; No trained embeddings: random TF and cell type vectors. **(c)** t-SNE visualization of the TF embedding vectors. Orange circles indicate related TFs that are in close proximity in t-SNE projection space: solid circles illustrate TFs from the same protein family, and dashed circles illustrate TFs having similar functions.

We additionally swap either or both types of embedding vectors during training to evaluate the contribution of the non-condition-specific network component. Replacing both types of embedding vectors during training results in a 26.2% drop in the mean auPRC score across all training conditions (Fig 4b). However, the significant performance decrease is mainly due to the removal of TF embeddings – separately removing TF embeddings and cell type embeddings result in a 25.7% and a 0.5% drop in the mean auPRC, respectively. Under the current model input feature setting, TF identity can only be learned through the TF embedding vectors. In contrast, cell-type-specific chromatin landscape can be learn from the cell-type-specific input features in addition to cell type embeddings. In the presence of cell-type-specific input features, cell type embeddings are used by the model to capture residual cell-type-specific information, and therefore only introduce marginal performance improvement (Figures 4 and S4).

Visualizing the trained TF embedding vectors in two dimensions using t-SNE [53] reveals that a subset of embedding vectors also reflects the TF functional similarities. Some TFs that are in close proximity in t-SNE space are from the same TF families, including FOXA1 and FOXA2, HNF4A and HNF4G, ATF2, ATF3 and ATF7, and JUN, JUNB, and JUND (Figure 4c, solid circles). Functionally related TFs such as IRF3 and STAT3 [54] are also adjacent to each other in t-SNE space (Figure 4c, dashed circle). However, these TF embedding vectors are explicitly trained to learn the biases introduced by TF labels. Available data for TFs of the same protein family are not necessarily from the same set of cell types. As a result, not all functionally related TFs are close in the t-SNE space, such as IRF (IRF3, IRF4, and IRF5) family proteins and TFs associated with c-Myc proteins (MAX and MAZ).

### 3.4. TF and cell type embeddings allow more reliable transfer predictions

Transfer learning allows models to make cross-TF and cross-cell-type predictions beyond training conditions. Existing single-task learners such as Catchitt achieve transfer learning by providing input features from a new cell type to a model trained on a different cell type. If multiple trained cell types are available for the same TF, the final cross-cell-type predictions are generated by averaging predictions from all trained cell types (Figure 5a, Average Train). A different transfer learning strategy proposed by AgentBind [25] involves pretraining a multi-TF model that does not disgintuish different TF identities before fine-tuning the model on a single TF of interest (No Embedding Transfer, Figure 5a). The former strategy cannot take advantage of the additional information introduced by other functionally related TFs, whereas the latter does not distinguish different TF identities in the multitask pretraining step. Since TF binding prediction can benefit from the multitask learning paradigm (Section 3.1), and a multitask learning model performance is highly influenced by the TF identity (Section 3.3), we hypothesize that NetTIME’s transfer learning strategy (Figure 5a, Embedding Transfer) is superior for cross-TF and cross-cell type binding prediction.

**Figure 5:**
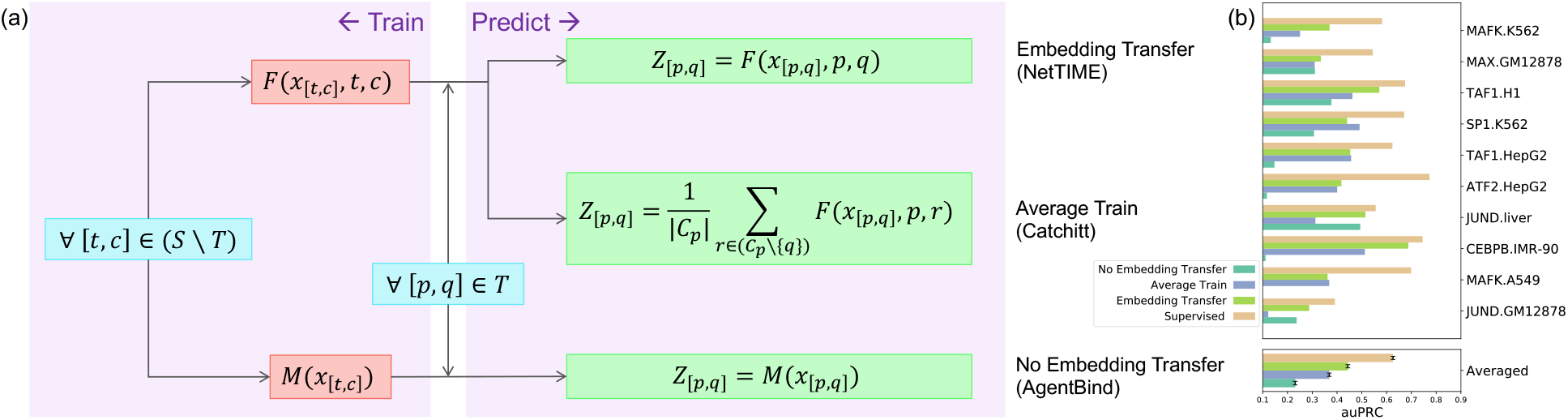
Comparison of different transfer learning strategies. **(a)** Detailed overview of the training scheme and prediction generation procedure using three transfer learning strategies implemented by NetTIME, Catchitt and AgentBind. [*t, c*] and [*p, q*] denote the particular TF (*t* or *p*) and cell type (*c* or *q*) combinations. *S* refers to the set of all conditions in our training dataset. *T* is the set of 10 leave-out conditions. *C*_*p*_ is the set of cell types that satisfy ∀*r* ∈ *C*_*p*_, [*p, r*] ∈ *S. x*_[*t,c*]_ is the input data from [*t, c*]. *Z*_[*p,q*]_ is the per-base-pair binding probability predictions for [*p, q*]. *F* and *M* can be any machine learning models. To avoid performance differences introduced by the model architecture, we use NetTIME model with TF and cell type embeddings (*F*) and with no embedding (*M*). **(b)** Transfer predictions using different transfer learning strategies are generated for 10 leave-out TF and cell type combinations within the training panels of TFs and cell types.

To evaluate the prediction quality of these three approaches, we pretrain a NetTIME model by leaving out 10 conditions for transfer learning. Transfer learning predictions are generally less accurate compared to supervised predictions (Supervised). For each transfer condition [*p, q*], we use the pretrained model to directly derive predictions for each transfer learning strategy (Figure 5a). However, transfer predictions generated by Embedding Transfer still significantly outperform those of the Average Train and the No Embedding Transfer (Figure 5b). Transfer predictions derived from NetTIME also achieve considerably higher accuracy compared to those from Catchitt and Leopard (Figure S5a and S5c). We additionally investigate whether different transfer learning strategies can benefit from fine-tuning by fine tuning all models using all conditions from TF *p* excluding [*p, q*]. This fine-tuning step additionally improves performance for Embedding and No Embedding Transfer approaches, whereas Average Train performance after fine-tuning remains low compared to two other approaches (Figure S5b and S5d).

Using trained TF and cell type embeddings additionally allows models to perform binding predictions beyond the training panels of TFs and cell types. We therefore test our model’s robustness when making predictions on unknown conditions using 4 conditions from 4 new TFs in 3 new cell types. Starting from a NetTIME model pretrained on all original training conditions (Section 2.1), we fine-tune the pretrained model for each transfer condition [*p*′, *q*′] separately by collecting available ENCODE datasets from all conditions from TF *p*′ and all conditions in cell type *q*′ excluding [*p*′, *q*′]. Transfer predictions generated from models trained with TF and cell type embeddings (Trained Embedding Transfer) significantly outperform those from models trained with no embeddings (No Embedding Transfer) that cannot distinguish different TF and cell-type identities (Figure 6a). TF binding motifs derived from predicted binding sites also show a strong resemblance to those derived from conserved ChIP-seq peaks (Figure 6b).

**Figure 6:**
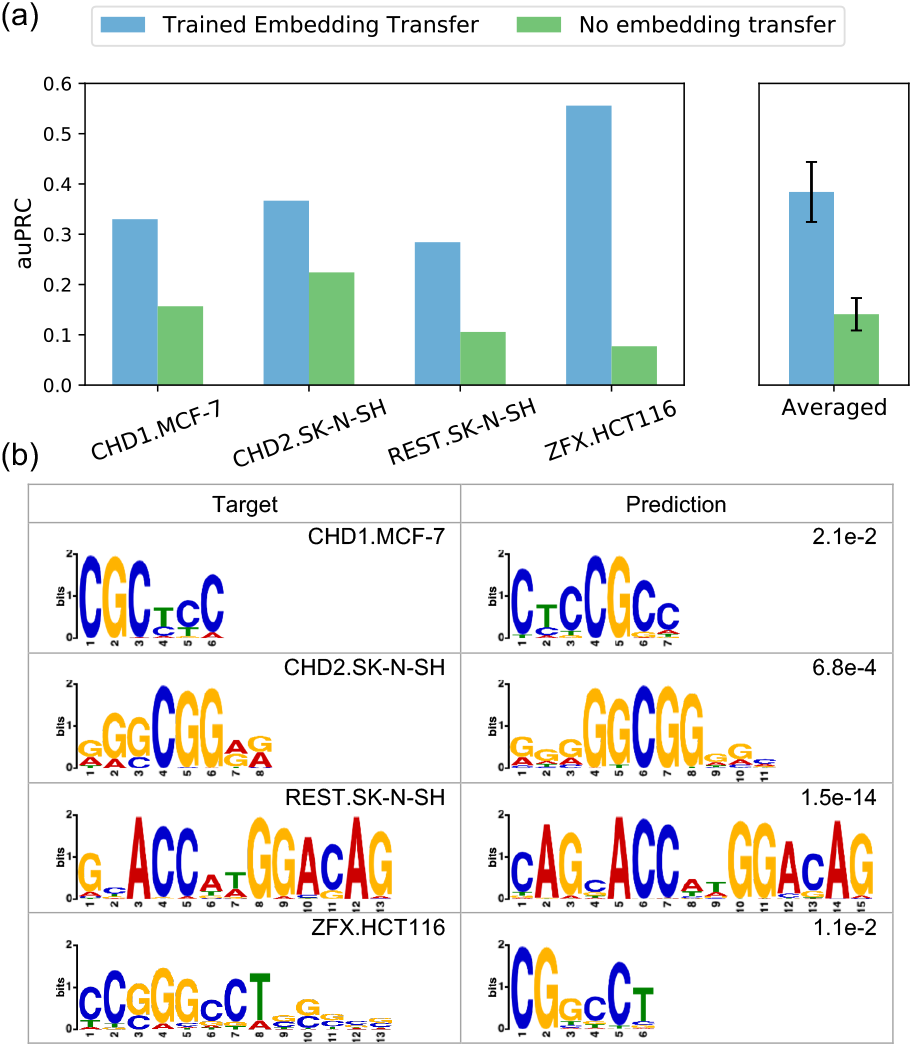
Transfer predictions to new TF and cell type combinations beyond the training panels of TFs and cell types. **(a)** Transfer predictions using models trained with either TF and cell type embedding vectors (Trained Embedding Transfer) or no trained embeddings (No embedding transfer). **(b)** Comparison of predicted TF binding motifs to those derived from target ChIP-seq conserved peaks. Predicted motifs are derived from Trained Embedding Transfer predictions. *De novo* motif discovery is conducted using STREME [55] software. Motif similarity p-values shown in the top right corner of the Prediction column are derived by comparing predicted and target motifs using TOMTOM [56].

### 3.5. A CRF classifier post-processing step effectively reduces prediction noise

Summarizing the binding strength, or probability, along the chromosome at each discrete binding site is an important step for several downstream tasks ranging from visualization to validation. Deriving binary binding decision from binding probabilities are typically done by finding a probability threshold that achieves the best prediction accuracy [35, 47, 11]. We test this baseline approach by evaluating the model’s predictive performance at 300 randomly selected probability thresholds. We find that at threshold 0.1302, the model achieves the highest IOU score of 36% (Figure 7a).

**Figure 7:**
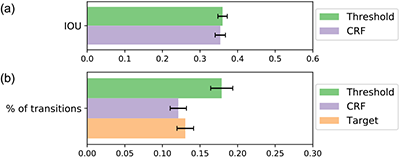
Binary classification performance using the probability threshold and CRF. Performance evaluated by **(a)** the mean IOU score and **(b)** the percentage of class label transitions per sequence (bottom), both calculated over all training conditions.

We alternatively train a CRF classifier, as a manually selected probability threshold is poorly generalizable to unknown datasets. These two approaches achieve comparable predictive performance as evaluated by IOU scores (Figure 7a). However, prediction noises manifested as high probability spikes are likely to be classified as bound using the probability threshold approach. To evaluate the effectiveness of reducing prediction noises using the probability threshold and the CRF approaches, we calculate the percentage of class label transitions per sequence within the target labels and within each of the predicted labels generated by these two approaches. The transition percentage using CRF is comparable to that of the true target labels, and is also significantly lower than the percentage obtained using the probability threshold approach (Figure 7b). This indicates that CRF is more effective at reducing prediction noise, and therefore CRF predictions exhibit a higher degree of resemblance to target labels.

## 4. Conclusions

In this work, we address several challenges facing existing methods for TF binding site predictions by introducing a multitask learning framework, called NetTIME, that learns base-pair resolution TF binding sites using embeddings. We show that our multitask learning approach improves prediction accuracy by increasing the data available to the model. Both the condition-specific and non-condition-specific components in our multitask framework are important for making accurate condition-specific binding predictions. The use of TF and cell type embedding vectors additionally allows us to make accurate transfer learning predictions within and beyond the training panels of TFs and cell types. Our method also significantly outperforms previous methods under both supervised and transfer learning settings, including Catchitt and Leopard.

Although DNA sequencing currently can achieve base-pair resolution, the resolution of ChIP-seq data is still limited by the size of DNA fragments obtained through random clipping. A considerable fraction of the fragments are therefore false positives, whereas many transient and low-affinity binding sites are missed [57]. Additionally, ChIP-seq requires suitable antibodies for proteins of interest, which can be difficult to obtain for rare cell types and TFs. Alternative assays have been proposed to improve data resolution [58, 59, 60] as well as to eliminate the requirement for antibodies [61, 62]. However, datasets generated from these techniques are rare or missing in data consortiums such as ENCODE [21] and ReMap [22]. NetTIME can potentially provide base-pair resolution solutions to more complex DNA sequence problems as labels generated from these alternative assays become more widely available in the future.

ATAC-seq (Assay for Transposase-Accessible Chromatin using sequencing [63]) has overtaken DNase-seq as the preferred assay to profile chromatin accessibility, as it requires fewer steps and input materials. However, these two techniques each offer unique insights into the cell-type-specific chromatin states [64], and it is therefore potentially beneficial to incorporate both data types for TF binding predictions. In fact, extensive feature engineering has been the focus of many recent *in vivo* TF binding prediction methods [52, 29, 20]. It is also important to note that, without strategies for handling missing features, increasing feature requirements significantly restricts models’ scope of application (Figure S1). A comprehensive evaluation of data imputation methods [65, 66, 67, 68] can be difficult due to the lack of knowledge of the true underlying data distribution. We plan to extend our model’s ability to learn from a more diverse set of features, and investigate more efficient ways to handle missing data. We also plan to explore other neural network architectures to improve model performance while reducing the model’s feature requirement.

NetTIME is extensible, and can be adapted to improve solutions to other biology problems, such as transcriptional regulatory network (TRN) inference. TRN inference identifies genome-wide functional regulations of gene expressions by TFs. TFs control the expression patterns of target genes by first binding to regions containing promoters, distal enhancers and/or other regulatory elements. However, functional interactions between TFs and target genes are further complicated by TF concentrations and co-occurrence of other TFs. A series of methods have been proposed for inferring TRNs from gene expression data and prior knowledge of the network structure [69, 70, 71]. Prior knowledge can be obtained by identifying open chromatin regions close to gene bodies that are also enriched with TF motifs [72]. However, this method is problematic for identifying TF functional regulations towards distal enhancers and binding sites without motif enrichment. *In vivo* predictions of TF binding profiles, however, can serve as a more flexible approach to generating prior network structure as it bypasses the aforementioned unnecessary constraints. In future work, we hope to adapt the NetTIME framework to explore more efficient approaches for generating prior knowledge for more biophysically motivated TRN inference.

## Acknowledgements

We thank members of the Bonneau and Cho labs for providing helpful suggestions on the project and on the manuscript. We thank the NYU High Performance Computing team and the Flatiron Institute Scientific Computing team for their excellent high performance computing support.

## Data Availability

The data used in this manuscript are downloaded from the ENCODE project website: https://www.encodeproject.org/. The assession numbers are provided in Supplementary Data 1-4. The ENCODE-DREAM Challenge data are downloaded from the Challenge website[4].

## Footnotes

1 http://dreamchallenges.org/about-dream/

2 https://www.encodeproject.org/about/experiment-guidelines

