## supplementary information for "NetTIME: a Multitask and Base-pair Resolution Framework for Improved Transcription Factor Binding Site Prediction"

*To whom correspondence should be addressed.

### 1 Supplementary Method

#### 1.1 Data retrieval and preprocessing

The list of 71 TF-focused ChIP-seq experiments we use to generate target labels are provided in Supplementary Data 1. Combined peak set are first generated by merging the conserved and relaxed peak sets from the above experiments. Peaks that are longer than 1000 bp are removed. Two overlapping peaks are merged when 1) they overlap for more than 200 bp, and 2) the resulting merged peak is shorter than 600 bp. Each interval in the merged peak set is used to create one 1000 bp example sequence where the midpoints of the example and the interval are the same. For TF $p$ in cell type $q$, a nucleotide $n$ is classified as bound if $n$ is within any ChIP-seq peaks for condition $[p, q]$, and unbound otherwise.

The ENCODE experiments used to generate cell type-specific features are provided in Supplementary Data 2. We download the narrowPeak BED files for all DNase-seq and histone ChIP-seq experiments, read-depth normalized signal bigWig files for all DNase-seq experiments, and the signal p-value bigWig files for all histone ChIP-seq experiment from the ENCODE Consortium. The genomic interval of each example sequence is intersected with DNase-seq and Histone ChIP-seq bigWig files to retrieve the corresponding example cell-type-specific feature signals. To reduce noise, only positive signals that fall within peak regions, defined in the DNase-seq and histone ChIP-seq narrowPeak BED files, are retrieved to generate the example feature signal tracks. Each cell-type-specific feature signal track across all examples are further zscore normalized before being used as input to the NetTIME model.

To test our model's ability to make transfer predictions beyond the training panels of TFs and cell types, we additionally collect target labels from 40 ENCODE TF-focused ChIP-seq experiments (Supplementary Data 3) and cell type-specific features from 10 cell types (Supplementary Data 4). These additional datasets are processed using the same procedures described above to generate additional fine-tuning examples.

#### 1.2 Training strategy and performance evaluation

NetTIME can be trained either in the supervised learning fashion or the transfer learning fashion. Training and validation data always contain the same set of conditions (i.e., TF-cell-type pairs). When evaluating model's supervised prediction performance, the test data contains the same set of conditions as the training and validation data. However, a new set of conditions unseen during training and validation is used to evaluate model's transfer learning performance at test time. NetTIME is implemented using PyTorch (v1.8.1) (Paszke, et al., 2019). We use the Adam (Kingma & Ba, 2014) optimizer with a batch size of 1800 to perform parameter update. learning rate is set to 1e-4 for the supervised learning experiments and the pretraining of the transfer learning experiments. Learning rate for transfer learning fine-tuning experiments is reduced to 5e-5. Models trained with different configurations are initialized using the same random seed to ensure the results are comparable and reproducible. Performance scores are calculated for each condition separately. P-values are derived from the one-sided Wilcoxon signed-rank test using performance scores derived from all conditions.

For each transfer learning experiment, we first define a set of conditions for which we intend to generate transfer learning predictions. At the pretraining stage, we train one supervised model by leaving out the transfer learning conditions. During fine tuning, we train separate models for each transfer learning condition (i.e., TF $p$ in cell type $q$) using all data belongs to TF $p$ and all data belongs to cell type $q$. Model's transfer learning performance is evaluated using test data from TF $p$ in cell type $q$.

The NetTIME neural network and CRF are trained sequentially. Predictions generated from the best-performing neural network checkpoint are used to train one CRF model across all conditions in the training set. Softmax-transformed neural network predictions are further log-transformed before given to the CRF model as input (Figure S2, (Richard & Lippmann, 1991)). The CRF model is implemented using the pytorch-crf (v0.7.2)[1]. We use the Adam (Kingma & Ba, 2014) optimizer with a learning rate of 1e-4 and a batch size of 2700 to optimize the model. The CRF checkpoint that achieves the lowest negative conditional log likelihood loss defined in Equation (4) is selected as the best CRF checkpoint.

### 1.3 Method comparison

#### 1.3.1 Regulatory-region-focused training and evaluation

The NetTIME model performance is evaluated on the sample level (the presence of TF binding event given a 1kb genomic sequence) and on the base-pair level using our test examples generated from chromosomes 1, 8 and 21. NetTIME's sample level performance is compared with Catchitt and Leopard, whereas its base-pair level performance is compared against Leopard. Sample-level predictions from NetTIME are generated by taking the maximum prediction score across all 1000-bp interval as well as the center 200-bp region for each example sequence in our test data. For both baseline methods, training, validation, and test data are split the same way as NetTIME (Table S1). Performance achieved on the test dataset is reported. Below are detailed procedures for generating predictions from the two baseline methods.

**Catchitt** We separately train 71 Cachitt models for all conditions we included in our training set. These models are trained using data generated from the whole genome following the Catchitt documentation[2]. We use the default setting, including b=50 and i=5, to train all models. For each condition, the DNase-seq bigwig file and the TF ChIP-seq conserved peak bed file used by NetTIME are provided to Catchitt to generate input features and target labels, respectively. The TF motif PWMs required by Catchitt are downloaded from HOCOMOCO (v11) motif database[3]. For each of the 1000-bp example sequence in our test data, final prediction score is calculated by taking the maximum prediction score from all intervals that overlap with the example sequence. The same procedure is used on the center 200-bp region for each example to generate the final prediction score at 200-bp resolution. To make transfer predictions using Catchitt, we provide input features belonging to the transfer condition to models trained in different cell types for the same TF. We further average predictions generated from multiple cell types of the same TF to derive final prediction scores for the transfer condition. The same procedure is also used by Catchitt to generate transfer predictions (Keilwagen, Posch, & Grau, 2019).

**Leopard** We separately train 71 Leopard models for all conditions we included in our training set. For each condition, we generate Leopard input features and target labels using the same DNase-seq bigwig file and the ENCODE conserved peak bed file as NetTIME. DNase-seq bigwig files across all cell types are quantile normalized before provided to Leopard. Each Leopard model is trained and validated using data from the same cell type with the default hyperparameter setting including lr=1e-3 and num_epoch=5. Leopard is a single-task learner that trains one model per TF and cell type. We therefore follow the same transfer prediction procedure as Catchitt to make transfer predictions using Leopard. Models trained (and validated) from different cell types of the same TF are used to derive base-pair level binding predictions by providing the models with input data from the transfer condition. These predictions are further averaged across all cell types of the same TF to derive the final target condition binding probabilities.

#### 1.3.2 Genome-wide training and evaluation using ENCODE-DREAM Challenge data

The ENCODE-DREAM Challenge data are directly downloaded from the Challenge website[4]. Train, validation and test data are split according to chromosomes in the same fashion as shown in Table S1. Methods are trained and evaluated in a within-cell-type and cross-chromosome fashion similar to the within-cell-type benchmarking round of the Challenge where models are trained, validated and tested on different chromosomes of the same TF and cell type combinations. We report model performance using 200-bp intervals from the test chromosomes provided by ENCODE. For NetTIME, the 1000-bp training examples are generated from the 200-bp ENCODE-DREAM intervals by expanding the window 400-bp on both

[1] https://pytorch-crf.readthedocs.io/en/stable/

[2] http://www.jstacs.de/index.php/Catchitt

[3] https://hocomoco11.autosome.org/downloads_v11

[4] https://www.synapse.org/\#!Synapse:syn6131484/wiki/402026

sides. We compute the maximum prediction score across the center 200-bp window to generate the sample-level prediction score for model performance evaluation. Catchitt and Leopard are trained and evaluated using the same set of hyperparameters mentioned in Section S1.3.1.

### 2 Supplementary Tables

| Dataset | Chromosomes | Peak set(s) | Number of conditions | Number of samples | |
|---|---|---|---|---|---|
| | | | | Per condition | Total |
| Training | 3-7, 10-20 | Conserved + Relaxed | 71 | 937,676 | 66,574,996 |
| Validation | 2, 9, 22 | Conserved | 71 | 101,644 | 7,216,724 |
| Test | 1, 8, 21 | Conserved | 71 | 102,178 | 7,254,638 |

**Table S1. The number of samples in training, validation and test datasets used to train and evaluate NetTIME performance.** Data splits are performed according to chromosomes. Both the ENCODE TF ChIP-seq conserved and relaxed peak sets are used for training, whereas only the conserved peak set is used to construct the validation and test datasets. A condition specifies a single TF-focused ChIP-seq experiment conducted in a particular cell type. All training, validation and test data are generated from the same set of 71 ChIP-seq conditions spanning 22 TFs and 7 cell types.

### 3 Supplementary Figures

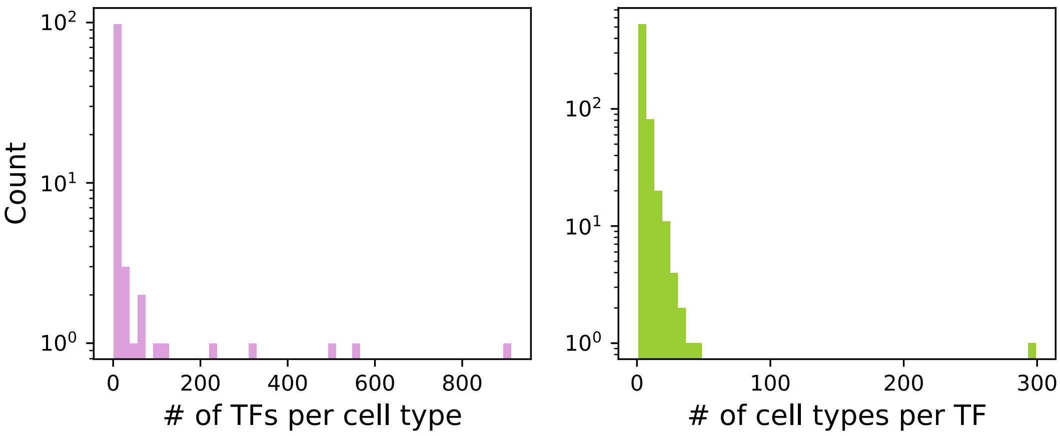


**Figure S1. ENCODE TF ChIP-seq experiments grouped by TFs and cell types.** Left: the number of TF ChIP-seq experiments per cell type. Right: the number of cell types in which a TF has ChIP-seq experiments. Both histograms skew towards the left, indicating that most cell types only have ChIP-seq data from a small number of TFs and most TFs only have ChIP-seq data in a small number of cell types. Data obtained from ENCODE Consortium[5] and reflects the database status as of December 2020.

[5] https://www.encodeproject.org/

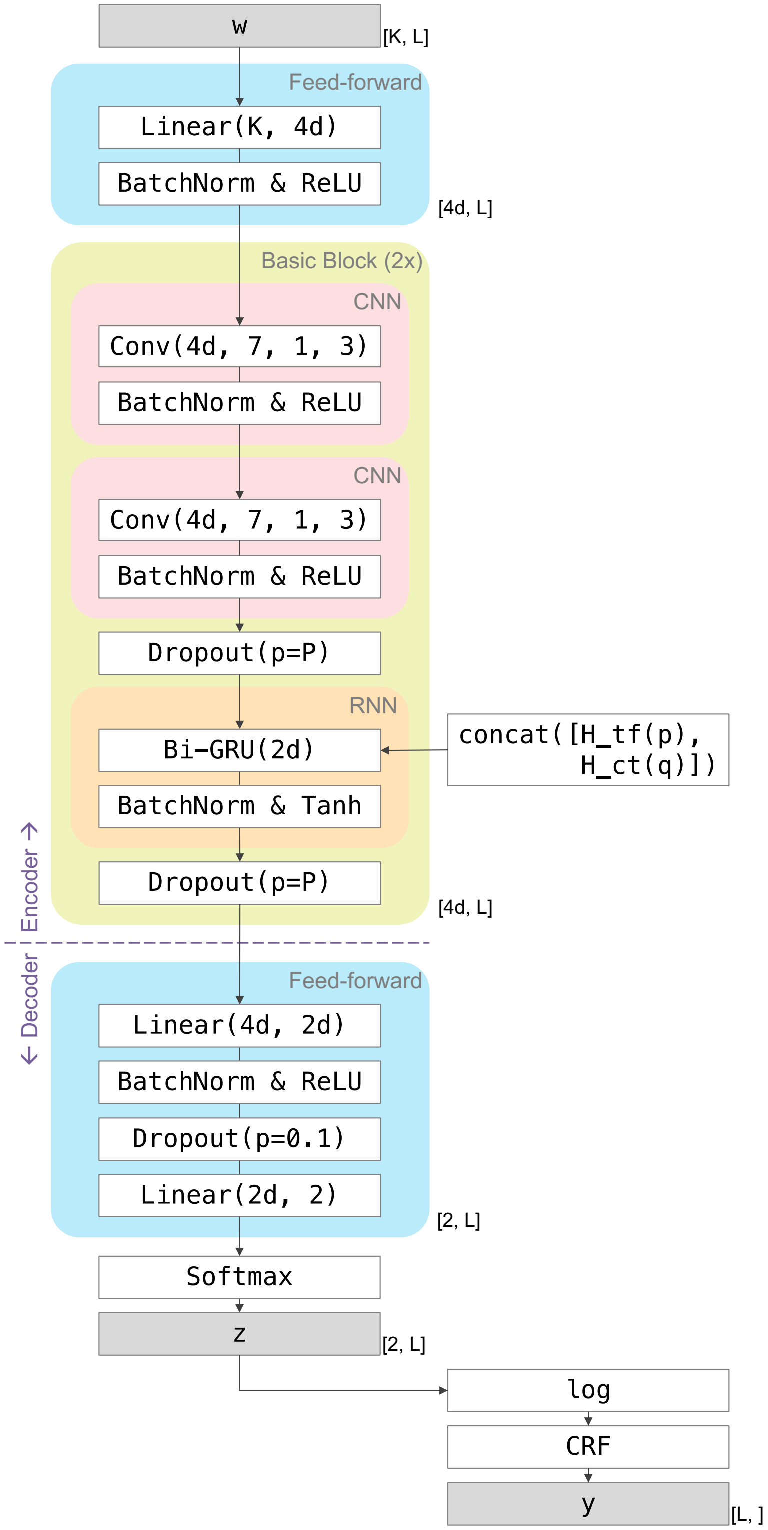


**Figure S2. NetTIME architecture.** Detailed view of the NetTIME architecture shown in Figure 1b. Three main parameters of the model are the number of input features $K$, example sequence length $L$, and embedding dimension $d$. Output dimensions for different model layers are shown on the right side of the architecture blocks. Linear$(a, b)$ denotes the linear transformation of input size $a$ and output size $b$; Conv$(e, f, g, h)$ denotes the 1D convolution operation of $e$ output channels, kernel size $f$, stride $g$ and padding $h$. Bi-GRU$(i)$ denotes bi-directional GRU layer of hidden size $i$. We set dropout probability $P = 0.1$ for the first pass of the Basic Block, and $P = 0.0$ for the second pass.

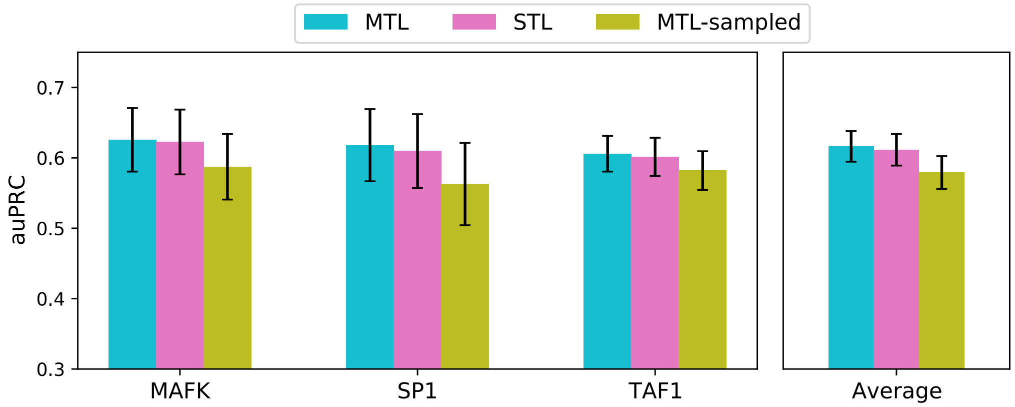


**Figure S3. Performance comparison between multitask learning and single-task learning approaches using three functionally unrelated TFs.** Models are trained with data from MAFK, SP1 and TAF1 across multiple cell types. Left panel shows the average performance measured by auPRC across multiple cell types of the same TF. Right panel averages performance across multiple TFs shown in the left panel. We observe marginal benefit (0.5% mean auPRC improvement) to using MTL strategy over STL when training unrelated TFs. However, the average auPRC score achieved by MTL-sampled is noticeably lower compared to STL. This additional analysis suggests that the relatedness among jointly trained conditions plays an important role in determining the effectiveness of multitask learning models. Increased data availability improves model generalization, although too much data heterogeneity reduces model predictability.

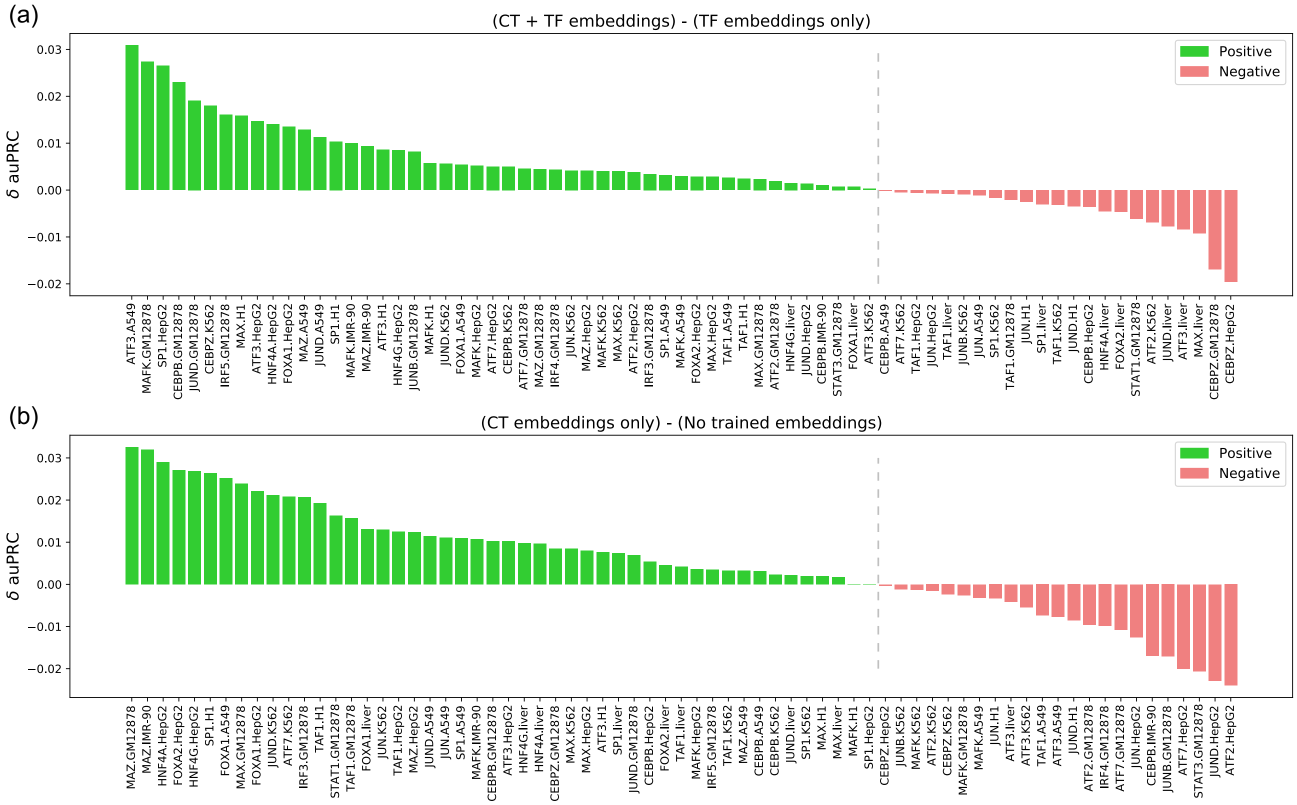


**Figure S4. Contribution of cell type embeddings during model training.** Detailed visualization of the auPRC score differences ($\delta$auPRC) for all 71 conditions **(a)** between CT + TF embeddings and TF embeddings only, and **(b)** between CT embeddings only and No trained embeddings. In both **(a)** and **(b)**, models trained with cell type embeddings (CT + TF embeddings and CT embeddings only) outperform their corresponding models trained without cell type embeddings (TF embeddings only and No trained embeddings) in 48 out of 71 conditions.

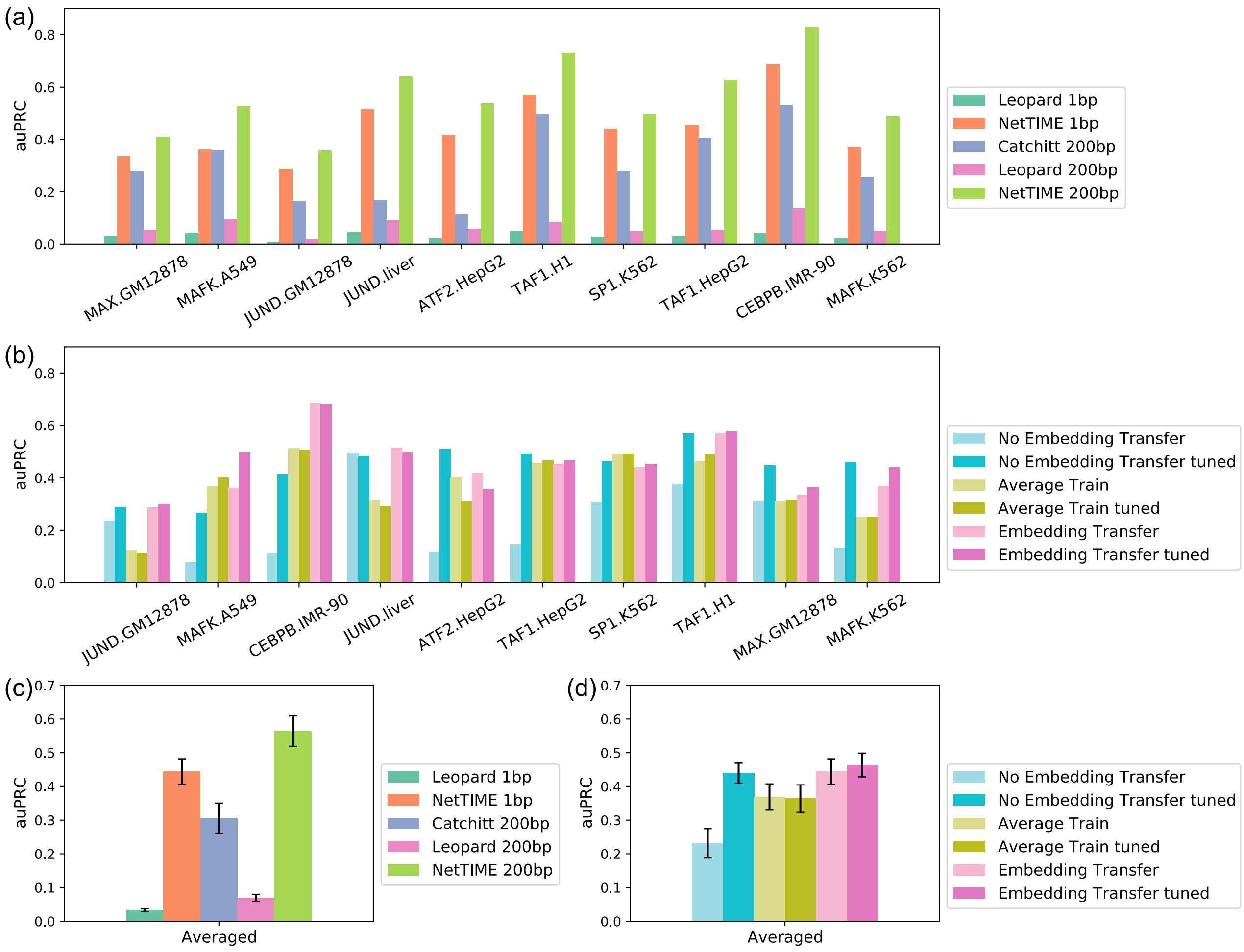


**Figure S5. Comparison of transfer learning strategies using 10 leave-out conditions within the training panels of TFs and cell types.** **(a)** and **(c)**: comparing transfer learning accuracy between NetTIME's Embedding Transfer approach with two other baseline methods under 1-bp and 200-bp resolutions. **(c)** is the mean auPRC score across conditions shown in **(a)**. Transfer predictions for Catchitt and for Leopard are achieved using the Average Trained method, where input features for the transfer conditions are given to models trained on different cell types of the same TF. The final prediction probabilities are derived by averaging all models' predictions if multiple trained models exist. Transfer predictions derived from both Catchitt and Leopard are significantly less accurate compared to NetTIME. **(b)** and **(d)**: transfer learning accuracy in pretrained and fine-tuned (tuned) NetTIME models using three transfer learning strategies. **(d)** is the mean auPRC score across conditions shown in **(b)**. The fine-tuning step improves the mean auPRC scores across all transfer conditions for both the Embedding Transfer and the No Embedding Transfer approaches. However, no improvement is observed after fine-tuning the Average Trained approach. Embedding Transfer outperforms two other approaches both with and without the fine-tuning step. More specifically, Embedding Transfer tuned outperforms the Average Train tuned and the No Embedding Transfer tuned by 9.4% and 2.4% on average, respectively. However, the benefit of having TF embeddings to distinguish TF identities is reduced after fine-tuning on data from the same TF, and we are able to observe significant improvements on the No Embedding Transfer model performance after fine-tuning.
